# Impact of land-use intensity, productivity, and aboveground richness on seed rain in temperate grasslands

**DOI:** 10.1101/2024.10.22.619565

**Authors:** Y.P. Klinger, S. Kunze, N. Hölzel, M. Freitag, V.H. Klaus, T. Kleinebecker, D. Prati, L Neuenkamp

## Abstract

1. Seed rain, the amount of seeds reaching an area via primary or secondary dispersal, affects the regeneration of plant communities and shapes the trajectory of future community composition. In agricultural grasslands, the composition and density in seed rain are mainly driven by land use, but drivers of seed rain quality and quantity along land-use gradients are poorly understood. We studied the effects of land-use intensity (LUI), its components (i.e. fertilization, mowing, and grazing), productivity, and aboveground vegetation composition and richness on seed rain.
2. We collected the seed rain over a five-month period in 142 grasslands and identified emerging seedlings. Grass seedlings dominated seed rain most likely due to their high abundance in vegetation and intense and early seed set. Only ten species accounted for approximately 80 % of seedlings, with grasses such as *Lolium perenne* and *Alopecurus pratensis* being most abundant. Forbs such as *Cerastium holosteoides* and *Veronica arvensis* were abundant in seed rain despite lower cover, probably due to early and prolonged flowering and high seed production.
3. Seed rain of grasses and forbs reacted differently to LUI and vegetation richness, LUI effects on grass seed rain mainly determined total seed density. Seed rain richness first increased with LUI, but decreased at higher LUI levels. Consequently, seed rain richness consistently increased with vegetation richness. This is reflected by a decrease in the abundance of stress strategists and an increase in ruderals in seed rain with increasing LUI and decreasing vegetation richness.
4. Among LUI components, fertilization intensity most strongly affected seed rain density and composition, with negative effects at intermediate fertilization intensities. Mowing once a year increased seed rain density, whereas it decreased at higher mowing frequencies. Grazing intensity reduced overall seed density and richness by reducing grass seed density, while forb seed density remained unaffected.

*Synthesis*: Land-use intensity and aboveground productivity significantly influence the species composition and seed densities in the seed rain of temperate agricultural grasslands. Higher land-use intensity and productivity increased seed production but reduced taxonomic and functional diversity in seed rain and may negatively impact ecosystem stability and resilience.

## Introduction

Seed rain, the amount of seeds reaching an area by means of primary or secondary dispersal, is a key process in plant community assembly that occurs between seed formation and germination (Arruda et al., 2018; Carvalho et al., 2021). Consequently, seed rain may strongly influence the regeneration potential, seed bank formation and population dynamics of plant communities (Auffret & Cousins, 2011). As seed rain results from sexual reproduction, it supports the maintenance of genetic diversity and increases the probability of species and communities to adapt to changing environmental conditions (Chung et al., 2023). At ecosystem level, seeds provide an important nutritional source for higher trophical levels such as beetles, birds and small rodents (Edwards & Crawley, 1999). Thus, knowledge about the composition of seed rain provides valuable insights into the reproductive potential of plant communities (Jefferson & Usher, 1989; Poschlod & Jackel, 1993).

The composition of seed rain may be particularly important in grassland communities (Page et al., 2002; Poschlod & Jackel, 1993), as these ecosystems typically consist of plants with low dispersal distances and many grassland species do not form persistent soil seed banks (Klaus, Hoever, et al., 2018; Klaus, Schäfer, et al., 2018; Ludewig et al., 2021). Further, grasslands with dense and species-rich seed rain might be more resilient to plant species losses due to disturbances as it is more likely that plants re-stablish directly from seeds or a regularly replenished persistent soil seed bank (Arruda et al., 2021; Hannah, 2022). Grasslands with low density and diversity in seed rain may rely heavily on clonal growth to sustain populations, which could result in low (genetic) diversity and low regeneration and dispersal potential. Thus, regular seed inputs via seed rain can be considered crucial to sustain grassland populations in the long term.

In agricultural grasslands, land-use intensity is a key driver of many ecosystem processes (Allan et al., 2014) driving plant diversity and composition and may thus strongly determine the composition and intensity of seed rain. Permanent agricultural grasslands (i.e. grasslands that exist for ≥5 years without being included in a crop rotation) are mostly maintained by agricultural practices such as fertilization, mowing, and grazing. Increased nutrient availability through fertilization results increased productivity, which allows more frequent and earlier mowing on meadows and higher livestock densities on pastures. Increased productivity leads to increased competition for light and the replacement of small- and slow-growing species by tall and fast-growing ones (Hautier et al., 2009), which is reflected in compositional shifts in the seed rain. Regarding seed density, fertilization and productivity have been shown to increase seed production under greenhouse conditions (Manning et al., 2009), but field studies have shown diverse and species-specific reactions to fertilization and raised productivity (Burkle & Irwin, 2010; Smith et al., 1996).

Usually, increased mowing or grazing intensity follow fertilization. Both, mowing machinery and grazing animals can serve as dispersal vectors for a number of grassland species and thus contribute to the genetic and functional connectivity between grasslands (Klinger et al., 2021). However, with increasing mowing frequencies and higher livestock densities, sexual reproduction and seed rain can be impeded due to the removal of plant tissue and reproductive organs, while the ability of clonal spread may become more important. Concerning seed rain, this is expected to reduce overall seed density (Martínková et al., 2020; Plantureux et al., 2005) and may lead to a higher dissimilarity between the standing vegetation and the species that manage to produce seeds, thus driving future community development. This is especially true in intensively mown meadows where silage is produced with three to six cuts per year and mowing takes place well before the first flowering. Similar effects have been observed in intensively grazed cattle pastures, where grazing starts early in the year and is carried out throughout the season with high stocking densities (Mielke & Wohlers, 2019). Thus, high land-use intensities in grasslands can severely limit the production, shedding, and dispersal of seeds, thereby constraining the regeneration, spatial and temporal spread of plant populations, that may even lead to local extinctions (Ozinga et al., 2009; Plantureux et al., 2005).

Up to now, no large-scale study has investigated the impact of land-use intensity on seed rain. Zeiter et al. (2013) studied seed rain of hay meadows across twelve sites in Switzerland in relation to a productivity gradient. By installing seed traps at ground level, they found an increased seed density and aboveground species richness with increasing productivity, which was explained by an increased seed production due to higher nutrient availability. Similar results were also found by Scotton (2020), who hand-collected mature seeds in permanent grasslands in the Italian Pre-Alps. However, the generality of these findings is still unclear, as all existing studies investigated not more as a handful of sites across rather short land-use and productivity gradients. For example, at very high land-use-intensities, a decrease of seed density and diversity of seed rain could be expected, as only few species manage to reproduce sexually under these conditions. The similarity between the aboveground vegetation and the seed rain can give further valuable information on the future development of grassland communities, as a high similarity between vegetation and seed rain may indicate stable plant communities whereas a low similarity may indicate community shifts, or the importance of clonal regeneration. Finally, assessing not only the response of taxonomic but functional composition of seed rain to land-use intensity and shifts in aboveground vegetation can add more mechanistic insights underlying variation in seed rain (Mayfield et al., 2010), which are of utmost importance to take effective management actions preserving grassland biodiversity and functioning.

We studied seed rain in 142 permanent temperate grasslands across a long gradient of land-use intensity in terms of mowing, grazing, and fertilization. As land-use intensity affects plant community composition also indirectly via productivity (Socher et al., 2012), we investigated the effect of aboveground plant biomass on seed rain. Species richness of the local plant community is another factor of high significance for the composition of seed rain (Harper, 1978) that was included in the analysis. We assessed the size, taxonomic and functional composition of seed rain in agricultural grasslands in relation to land-use intensity, productivity and species richness of the established vegetation and hypothesized

i. A unimodal relationship between land-use intensity and the total amount of seeds in seed rain
ii. A positive relationship between productivity and the total amount of seeds in seed rain
iii. A negative relation between species richness in the seed rain and land-use intensity.
iv. An increasing similarity between seed rain and established vegetation with increasing land-use intensity.

In particular, we addressed the following questions:

a. What is the functional and species composition and diversity in the seed rain?
b. How do land-use intensity and its components (fertilization, mowing, grazing), aboveground productivity, and aboveground richness shape seed density, diversity, and functional composition in seed rain?
c. How do land-use intensity, aboveground productivity and aboveground richness shape the similarity between aboveground vegetation and seed rain?

## Materials and Methods

### Study Area

We collected seed rain in three German regions in the framework of the Biodiversity Exploratories (Fischer et al., 2010). The three regions cover gradients in soil characteristics, elevation and climate that are representative of large parts of Central Europe. The regions comprise i) the Biosphere Reserve Schwäbische Alb, a calcareous mid-mountain range (48.4°N, 9.4°E), ii) the National Park Hainich-Dün, a calcareous low-mountain range 51.1°N, 10.4°E) and iii) the Biosphere Reserve Schorfheide-Chorin, a postglacial landscape (53.0°N, 14.0°E). While grasslands in the Schwäbische Alb and Hainich-Dün are located exclusively on loamy mineral soils, grasslands in Schorfheide-Chorin are found on drained fen soils rich in organic carbon or slightly acidic sandy soils (Fischer et al., 2010). We studied approx. 50 permanent grasslands per region on 50 × 50 m plots representing the most common soil types and comparable and-use intensity gradients within each region (Fischer et al., 2010).

### Land-use intensity

The compound land-use index (LUI) by Blüthgen et al. (2012) is based on standardized information on mowing, grazing, and fertilization practices reported annually by the landowners (Vogt et al., 2019) and was calculated using the LUI calculation tool (Ostrowski et al., 2020), which is implemented in the Biodiversity-Exploratories Information System (BExIS; http://doi.org/10.17616/R32P9Q). Information about grassland management from 2016 to 2018 was used to quantify land-use intensity. The number of cuts, the number of livestock units per ha and year and the amount of nitrogen fertilizer per ha and year were standardized by dividing raw values by the regional mean separately for the years 2016 to 2018. The compound LUI was calculated as the square root of the sum of the three standardized single components. Low LUI corresponds to unfertilized sheep pastures or meadows cut once per year, while high LUI indicates highly fertilized grasslands with high cutting frequencies and/or livestock densities.

### Vegetation surveys

Vegetation surveys were performed on a 4 m × 4 m subplot in all 150 grassland plots, all vascular plan species were identified and the percentage cover of each species was estimated. Surveys were done yearly from beginning of May to the beginning of June in 2016 to 2018 in all study regions. Simultaneously, aboveground plant biomass was harvested on each plot as a proxy of productivity. Plant biomass of eight 0.5 m × 0.5 m subplots was clipped at the height of 2 cm, dried at 80°C for 48 hours and weighed. The vegetation surveys and biomass data were averaged for the three surveyed years. One plot was excluded due to deviation from the biomass sampling protocol. This work is based on data elaborated by the Botany (core) project of the Biodiversity Exploratories programme (Bolliger et al. 2018, Fischer et al. 2017, Schäfer et al. 2018).

### Seed rain sampling

For seed rain collection, we installed five aluminum trays (13 cm × 10.3 cm × 4 cm; total area: 670 cm² per plot) filled with 300 cm² sterilized sand and drainage holes on each of the 142 grassland plots. Due to differences in plot layout, the trays were arranged in two different ways: On 75 plots, the trays were installed on the edges of the 50 m × 50 m plots (25 m distance between trays). On 71 plots, trays were installed on the edges of a 7 m × 7 m subplot within the 50 m × 50 m plot (3.5 m distance between trays). Testing for differences between the two setups with ANOVA revealed no significant differences in the data for total seed density and species richness in seed rain (S1 Figure 1).

**Figure 1:**
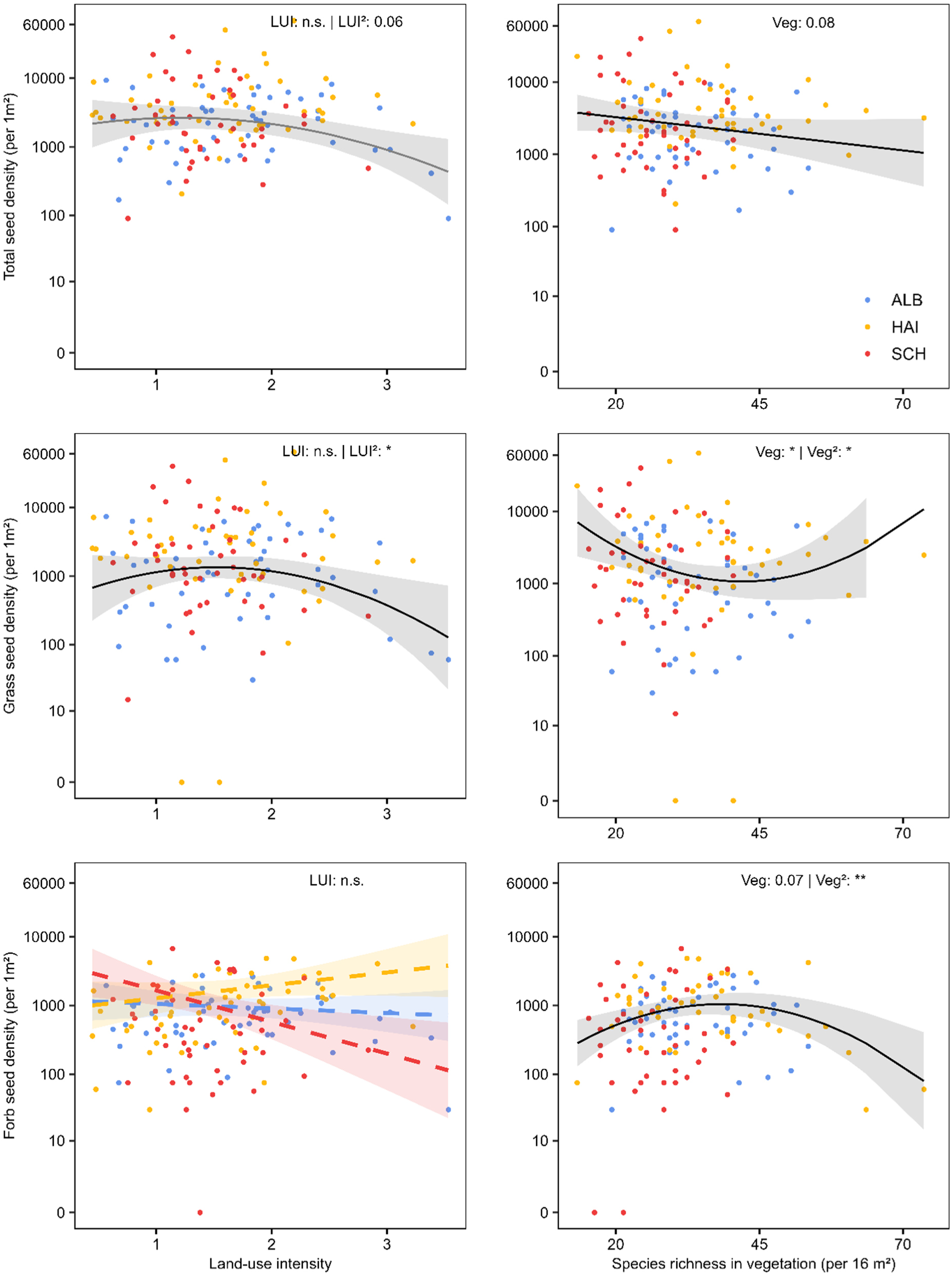
Effects of land-use intensity (LUI) and productivity on seed rain characteristics: (a-b) total seed density and seed density of different functional groups of (c-d) grasses and (e-f) forbs (including legumes). All y-axes were log-transformed. Predicted regression lines of the fixed effects based on the mean values of the other predictor variable and region Alb. Grey areas display the confidence interval (± 1.96 × SE) of the regression lines. The dots represent the observed densities and colours the study regions (Schwäbische Alb (ALB, blue), Hainich-Dün (HAI, orange) and Schorfheide-Chorin (SCH, red). Asterisks indicate significance levels: * p <0.05; ** p <0.01; *** p <0.001 and ‘n.s.’ = no significance. Marginally non-significant p values are recorded. See Supplementary material 2 for model results.

To collect seeds shed over the main vegetation period, we installed the trays between March 12 and April 14 and recollected them between August 4 and August 18, but few trays could only be recollected on September 26 when cattle grazing ended (114-195 days, on average 133 days). Of the 724 trays installed in the field, 109 trays (15 %) were damaged by cattle or land management and were excluded from the analysis (number of intact trays per study plot: 0 trays: 1 plot; 1 tray: 2 ×; 2 trays: 12 ×; 3 trays: 19 ×; 4 trays: 28 ×; 5 trays: 84 ×). Three plots with less than two remaining trays were removed before statistical analysis. Seed rain data is based on Hölzel et al. (2022b, 2022a)

For seedling counting and analysis, trays of each plot were pooled. From this data, we calculated seed density as the number of seeds per m² and species richness of seeds as the number of seedling species per m². To examine the pairwise dissimilarity between aboveground vegetation and seed rain in relation to the land-use intensity and productivity, we calculated the Sørensen dissimilarity index as a measure of beta diversity. The Sørensen index is defined /3sør = (b+c)/(2a+b+c) with a for species present in aboveground vegetation and in seed rain, b for the species present in vegetation but not in seed rain and c for the species present in seed rain but not in vegetation (Sørensen 1948).

All samples were stored in a refrigerator at 5°C for four weeks until the start of the greenhouse experiment. The amount of germinable seeds was then determined using the emergence method (Roberts, 1981). The sample material was pooled by plot (up to 1,500 cm³ if all trays were available) and spread on 20 cm × 30 cm styrofoam trays filled with sterile garden soil (Fruhstorfer Erde LD80 Archut^®^). In a greenhouse, these trays were exposed to controlled diurnally alternating temperatures (day: 18–24°C, night: 12–18°C), light (>10,000 lx from 6:00 a.m. to 10:00 p.m.), and humidity (<70%) conditions and were watered every three days. We added four control trays containing sterile garden soil to account for wind dispersed seeds in the greenhouse. Species germinating from these control trays (*Betula* spec. and *Juncus effusus*) were excluded from the analysis. To determine the species composition of the seed rain, we regularly counted and identified emerging seedlings to species level over seven months (September 2018 – April 2019). The soil was regularly disturbed using a fork to bring seeds to the surface (ca. twice per sample). After the fourth and fifth month, we fertilized the samples to stimulate germination in seeds that had not yet germinated. If seedlings were not identifiable at an early seedling stage, they were transplanted to pots and grown until identification was possible.

### Functional characteristics of seed rain

We investigated community-level traits relevant for the functional composition of seed rain. We calculated the total amount and community-weighted means (CWM) of seed mass of seed rain and the CWM of Grime’s strategy types (Grime, 1974) for the aboveground vegetation and seed rain. Seed weights for each species were derived from the LEDA database (Kleyer et al., 2008) by calculating the mean value per species, species missing from LEDA (two species) were derived from the Seed Information Database (ser-sid.org). Categorization and weighing of CSR strategy types for the calculation of CWM values was performed following Hunt et al. (2004), CSR values for species not listed there were derived from the Biolflor database (Kühn et al., 2004). Relative abundances for each species were calculated by setting the sum of all species abundances per vegetation relevée and the sum of all seed rain seedling numbers per plot, respectively.

### Statistical analysis

All statistical analyses were done with R 4.1.3 (R Core Team 2020) in the RStudio environment (RStudio 2021.09.0+351), graphs were created using the packages ‘ggeffects’ (Lüdecke, 2018) and ‘ggplot2’ (Wickham, 2016). For analyses, all numeric predictor variables were scaled to zero mean and unit standard deviation.

As response variables, we included the total seedling density and seedling density of grasses and forbs, seedling species richness, dissimilarity between vegetation and seed rain. As functional characteristics, we investigated, total seed CWM mass and the CWM abundance of C, S, and R strategists. The response variables were related to land-use intensity (LUI), productivity, and aboveground species richness. Soil pH_CaCl2_ (Schöning et al., 2021), soil C_org_ (Schöning, 2023), and the topographic wetness index (Magdon, 2024) were included as covariates as they comprise master variables characterizing local soil conditions and hence vegetation composition at each plot. For all response variables, a full model was defined using LUI, productivity, and aboveground richness as well as their interaction with region, and the covariates. Separate models were specified for the response variables seed density as well as for the density of grass and forb seeds. Because of the low number of legume seedlings (336), these were included to the forb seedlings for modelling.

Analyses were performed using generalized linear models (GLMs). In the GLMs, gaussian (function *glm)* or negative binomial models (function *glm.nb* from ‘MASS’ package; Ripley et al., 2024) were used for the density and total seed mass models, whereas for the dissimilarity and CWM models, beta family models with logit links from the glmmTMB package (Brooks et al. 2017) were used. The base function of all models was: response ∼ (LUI + productivity + aboveground species richness) * region + region + soil pH + topographic wetness index + soil C_org_. As environmental predictors can form quadratic i.e. hump-backed relationships with vegetation parameters (Fraser et al., 2015), we included LUI, productivity, and aboveground richness as linear and quadratic terms when a nonlinear relationship was evident and AIC (Akaike Information Criterion) decreased when adding a quadratic term to the model. Based on this model, model selection was carried out using the stepAIC function from the ‘MASS’ package (Ripley et al., 2024). Additionally, we fitted separate models for seed density (total, grasses, forbs) and species richness in seed rain in response to the LUI components mowing, grazing, and fertilizing. For this, we replaced LUI by mowing, grazing or fertilization intensity in the previous models but used the same error distribution and links as before.

Non-collinearity of all predictor variables was checked by Pearson’s correlation coefficient (thresholds ≥ 0.6). Significances were tested using type II (type III in the presence of significant interactions) likelihood ratio tests. All models were validated using the ‘DHARMa’ package (Hartig & Lohse, 2022). The explained variation by the models was derived using Nagelkerke’s pseudo-R². Since there is no calculation of pseudoR² for models fit by the R function *glmmTMB* from the glmmTMB package (Brooks et al. 2017) deriving the explained variation by the model, the squared correlation (cor²) between response and the predicted values was calculated.

## Results

### Functional and species composition in the seed rain

In total, 39,777 seeds of 126 vascular plant species germinated from the seed rain and we found a species pool of 366 species in the aboveground vegetation. Only nine species present in seed rain were not found in the aboveground vegetation. Mean seed density was 5,043 seedlings per m² (ranging from 90 to 68,589 seeds per m²) and mean species richness was 12.6 (ranging from 3 to 27 species). In total, we found 31,015 seeds (78 %) of 33 grass species, 8,413 seeds (21 %) of 82 forb species and 338 seeds (0.8 %) of ten legume species (Table 1). The study regions strongly differed in seed densities and species richness in the seed rain: The Hainich region had the highest seed density (7,109 ± 11,913 seeds m^-^²), whereas the Alb region was characterized by the lowest density (2,899 ± 2,415 seeds m^-2^), but had the highest mean species richness per seed rain sample (15; Table 1).

**Tab 1:** Total number of germinable seeds, mean values, standard deviation (SD), minimum (min) and maximum (max) values of seed density and species richness in seed rain samples (N = 142). Values shown cover all plots (total) and are separated for the functional groups of forbs, grasses, and legume, and also by study regions. Study regions: Alb = Schwäbische Alb, Hai = Hainich, Sch = Schorfheide.

| | Total seed count | Seed density ($m^{-2}$) | | Species richness (n) | |
| --- | --- | --- | --- | --- | --- |
| | | mean ± SD | min – max | mean ± SD | min – max |
| **All** | 39777 | $5043 \pm 8439$ | 90 – 68589 | $12.6 \pm 5.8$ | 3 – 27 |
| **Functional groups** |  |  |  |  |  |
| Grasses | 30990 | $3963 \pm 8082$ | 0 – 63824 | $6.6 \pm 3.5$ | 0 – 16 |
| Forbs | 8216 | $1014 \pm 1143$ | 0 – 6617 | $5.3 \pm 2.9$ | 0 – 15 |
| Legumes | 338 | $41 \pm 82$ | 0 – 473 | $0.7 \pm 1$ | 0 – 4 |
| **Study regions** |  |  |  |  |  |
| Alb | 7954 | $2899 \pm 2415$ | 90 – 9286 | $15.1 \pm 5.9$ | 3 – 27 |
| Hai | 21696 | $7109 \pm 11913$ | 205 – 68589 | $13.9 \pm 5.2$ | 5 – 25 |
| Sch | 10702 | $5132 \pm 7650$ | 90 – 39806 | $8.4 \pm 3.5$ | 3 – 17 |

The species with the highest number of seeds was *Poa trivialis* (12,750; 32 % of all seeds). The forb with the highest number of seeds was *Cerastium holosteoides* with 3,336 seed counts (8 %) and the legume with the highest number of seeds was *Trifolium repens* (113; 0.3 %). The 10 most frequent species emerging in our germination experiment comprised eight grasses (*Poa trivialis*, *Lolium perenne*, *Alopecurus pratensis*, *Dactylis glomerata*, *Poa pratensis agg.*, *Bromus hordeaceus agg.*, *Festuca rubra agg.*, *Holcus lanatus*) and two forbs (*Cerastium holosteoides*, *Veronica arvensis*), representing in total 79 % of all seedlings. A third (32 %) of all identified species occurred in less than four plots. *Cerastium holosteoides* was the most frequent species, which occurred in 109 of 142 plots (77 % of all plots).

### Land-use intensity effects on seed rain

Total seed density showed a hump-shaped distribution along the land-use intensity gradient (χ²_LUI_² = 3.6, p = 0.06; Supplementary material 2, Figure 1a). Seed density slightly increased to an optimum of LUI = 1.2 and then decreased for higher LUI values. Productivity and aboveground species richness had no significant effect on total seed density, but seed density tended to increase with increasing aboveground productivity (χ²_productivity_= 3.2, p = 0.07) and decreased with increasing aboveground richness (χ²_Aboveground richness_= 3.0, p = 0.08; Supplementary material 2, Figure 1b). The variation explained by the model including LUI, productivity, and aboveground richness was 28 % (Supplementary material 2). When modelling seed density separately for the different functional groups, grass seed density had a hump-shaped relation with LUI and aboveground richness. While grass seed density decreased above and below LUI = 1.5 (χ²_LUI²_ = 4.5, p < 0.05; Supplementary material 2, Figure 1c), the relation between grass seed density and aboveground richness was inverted, increasing above and below 42 species * 16 m^-^² (χ²_Aboveground richness_= 5.8, p < 0.05, χ²_Aboveground richness²_ = 6.2, p < 0.05). The variation explained by the model containing LUI, productivity, and aboveground richness was 11 % (Table 2). Forb seed density was not significantly affected by LUI (p = 0.5; Supplementary material 2, Figure 1e) or productivity (p =0.05), but showed a hump-shaped relationship with aboveground richness with an optimum at 38 species * 16 m^-^²(χ²_Aboveground richness_= 3.3, p = 0.07, χ²_Aboveground richness²_ = 8.8, p < 0.01; R² = 0.29; Supplementary material 2, Figure 1f).

Considering the effect of land-use intensity on species richness in seed rain, aboveground species richness also showed a hump-shaped relationship with LUI (χ²_LUI_² = 6.1, p < 0.05; Supplementary material 2, Figure 2a), no significant relationship with productivity (χ²_Productivity_ = 12.5, p < 0.001), and a positive relationship with plant species richness in aboveground vegetation (χ²_Aboveground richness_ = 3.9, p < 0.05; Supplementary material 2, Figure 2b). Regarding the LUI effects, seed species richness increased with LUI for LUI values up to 1.4 and decreased with higher LUI values. Species richness in seed rain differed among the regions in the order Alb > Hainich > Schorfheide. The variation explained by the model including LUI, productivity, and plant species richness in aboveground vegetation was 38 % (Supplementary material 2).

**Figure 2:**
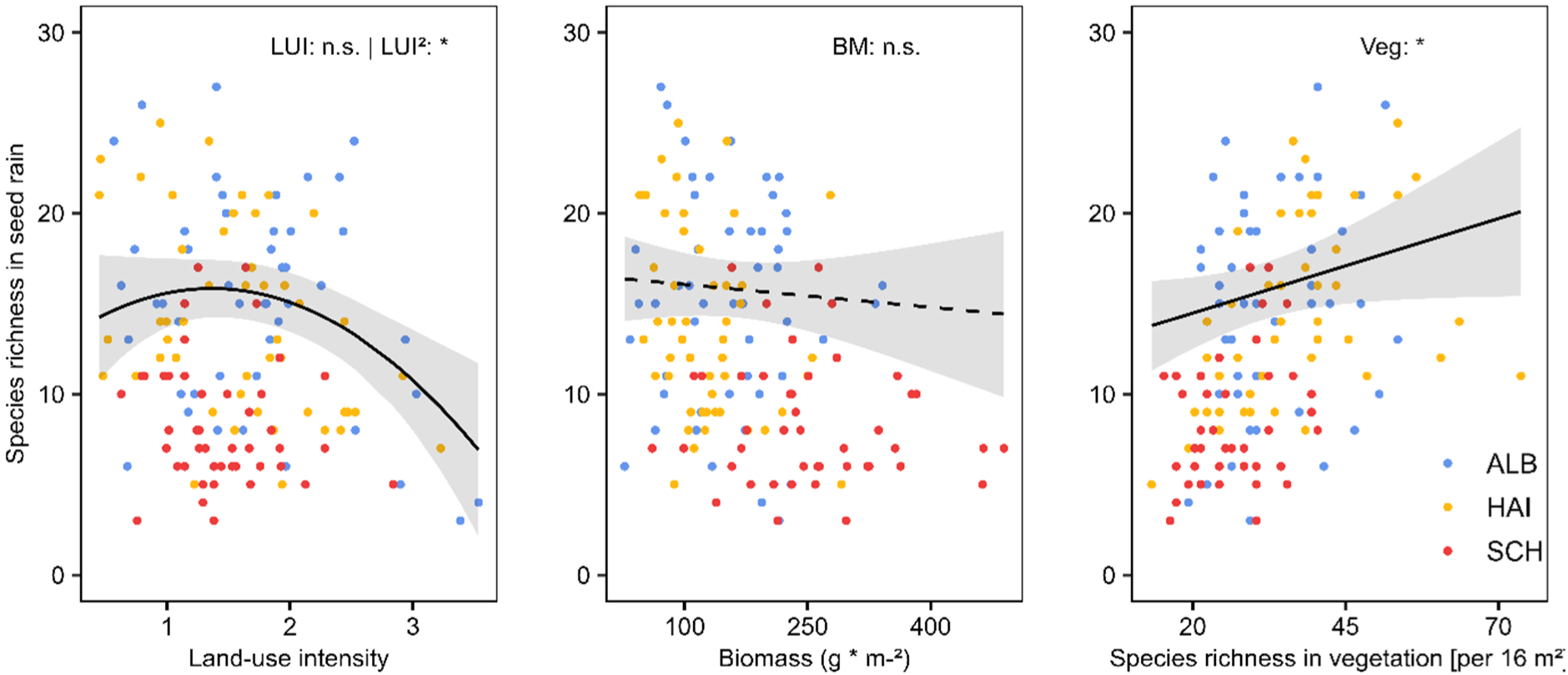
Effects of (a) land-use intensity (LUI) and (b) species richness in aboveground vegetation on species richness in seed rain. Predicted regression lines of the fixed effects based on the mean values of the other predictor variable and region Alb. Grey areas display the confidence interval (± 1.96 × SE) of the regression lines. The dots represent the observed species richness and colours the study regions (Schwäbische Alb (ALB, blue), Hainich-Dün (HAI, orange) and Schorfheide-Chorin (SCH, red). Asterisks indicate significance levels: * p <0.05; ** p <0.01; *** p <0.001 and ‘n.s.’ = no significance. See Supplementary material 2 for model results

Concerning the functional community characteristics of seed rain in relation to LUI, productivity, and aboveground richness, we found a hump-shaped relationship between the total seed mass in seed rain and LUI, where LUI stayed stable up until ∼1.2 and decreased afterwards (χ²_LUI_² = 5.2, p < 0.05, R²=0.47, Fig 4. a), whereas productivity and aboveground richness had no effect. The CWM seed mass increased linearly with increasing aboveground richness (χ²_Aboveground richness_ = 3.9, p < 0.05, Fig. 4b), whereas LUI had marginally non-significant effects (χ²_LUI_ = 3.8, p = 0.053, R²=0.37). Increasing LUI led to a significant decrease in the abundance of S strategists in seed rain (χ²_LUI_ = 15, p < 0.001, R² = 0.42) and resulted in an increased abundance of R strategists in seed rain (χ²_LUI_ = 12.5, p < 0.001, R² = 0.27). There was no effect of LUI, productivity or aboveground richness on the abundance of C strategists in seed rain.

**Figure 3:**
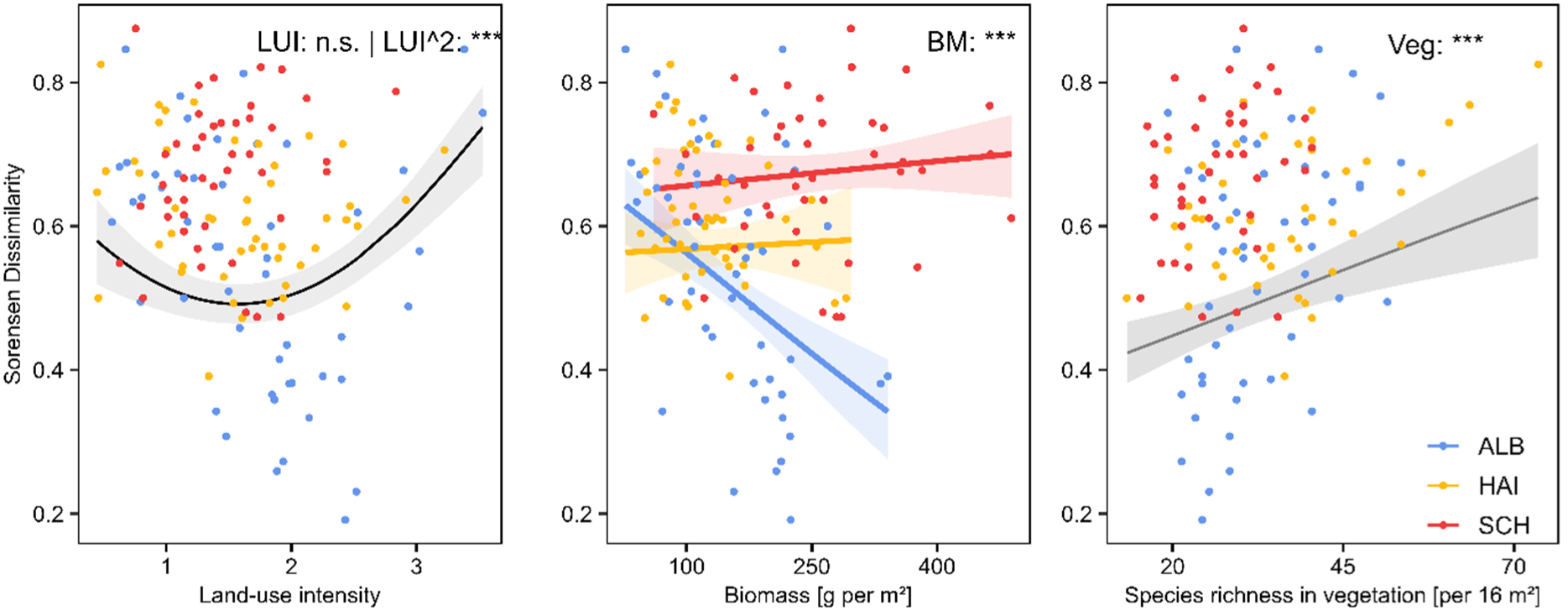
Effects of (a) land-use intensity (LUI) and (b) productivity (separately for each region due to a significant interaction between productivity and region), and (c) species richness in vegetation on the Sørensen dissimilarity index. Predicted regression lines of the fixed effects based on the mean values of the other predictor variable and region Alb. Grey areas display the confidence interval (± 1.96 × SE) of the regression lines. The dots represent the calculated dissimilarities and colors the study regions (Schwäbische Alb (ALB, blue), Hainich-Dün (HAI, orange) and Schorfheide-Chorin (SCH, red). Asterisks indicate significance levels: * p <0.05; ** p <0.01; *** p <0.001 and ‘n.s.’ = no significance. See Supplementary material 2 for model results.

**Figure 4:**
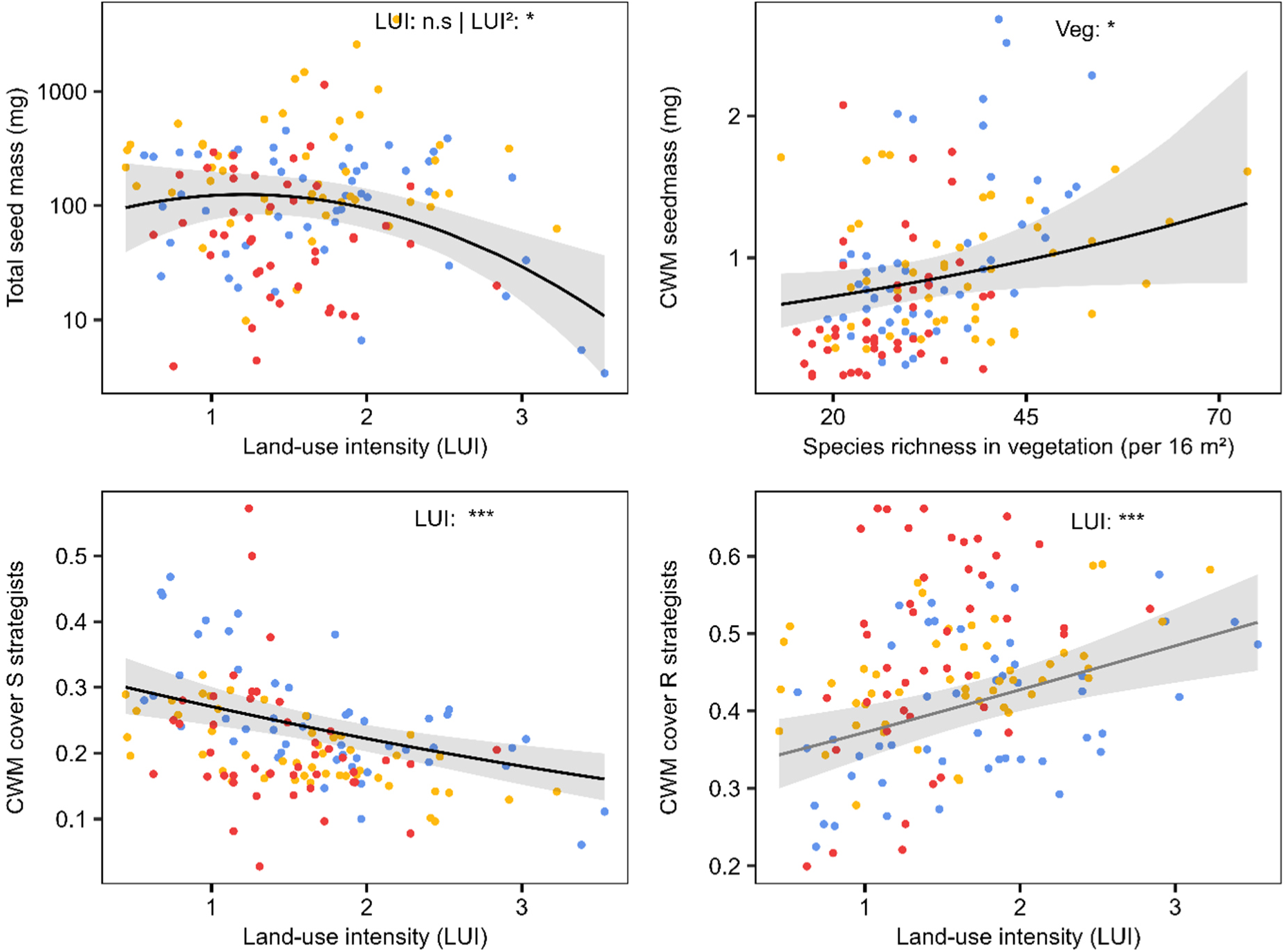
Effects of (a) land-use intensity (LUI) on total seed mass in seed rain, (b) species richness in vegetation on community-weighted mean (CWM) seed mass, (c) LUI on the CWM cover of S strategists, and (d) LUI effects on the CWM cover of R strategists. Predicted regression lines of the fixed effects based on the mean values of the other predictor variable and region Alb. Grey areas display the confidence interval (± 1.96 × SE) of the regression lines. The dots represent the calculated values per plot and colors the study regions (Schwäbische Alb (ALB, blue), Hainich-Dün (HAI, orange) and Schorfheide-Chorin (SCH, red). Asterisks indicate significance levels: * p <0.05; ** p <0.01; *** p <0.001 and ‘n.s.’ = no significance. See Supplementary material 2 for model results. Mind the different scales of y-axes.

Separate models on the effects of the single LUI components (fertilization, mowing, grazing) revealed consistent hump-shaped relationships of fertilization with total (χ²_Fertilization2_=6.85, p<0.01, optimum = 2.5, R²=0.41), grass (χ²_Fertilization2_=4.8, p<0.05, optimum = 2.5, R²=0.32) and forb seed density (χ²_Fertilization2_=6, p<0.05, optimum = 2.5, R²=0.33), as well as seed rain species richness (χ²_Fertilization2_ = 19.5, p<0.001, R²=0.72; Supplementary material 1, Figure S2). In particular, seed rain species richness increased at low to medium fertilization intensity of 2.7 and decreased afterwards (Figure S2). Mowing frequency showed similar hump-shaped relationships with total seed density and seed rain species richness having constant effects up to low to medium mowing frequencies of ∼ one cut per year and negative effects afterwards (Figure S1). Effects were strongest for species richness in seed rain (χ²_Mowing2_=6.3, p < 0.05, R²=0.42) and weaker and marginally non-significant for total seed density (χ²_Mowing2_= 3.3, p = 0.07, R²=0.3). Grass density decreased with increasing mowing frequency in the Hainich and Alb regions but increased in the Schorfheide region whereas forb seed density showed no significant response to mowing frequency. Grazing intensity linearly reduced seed richness (χ²_Grazing_=6.4, p<0.05, R²=0.64), and showed a hump shaped relation with grass seed density (χ²_Grazing²_=4.2, p<0.05, R²=0.27), with no effects on total and forb seed density (Figure S1).

### Effects land-use intensity on the compositional similarity between aboveground vegetation and seed rain

We found on average 28 % (± 13) of the plant species present in the aboveground vegetation also in the seed rain. The Sørensen dissimilarity index was significantly affected by land-use intensity (χ²_LUI_² = 27.3, p < 0.001) in a hump-shaped manner, decreasing up to a LUI of about 1.6, and increasing for higher values (Figure 3a). The effect of productivity on the dissimilarity between aboveground vegetation and seed rain varied between regions, with dissimilarity increasing with aboveground productivity in the regions Schorfheide and Hainich regions, but dissimilarity strongly decreasing with aboveground productivity in the region Alb (Figure 3b). Dissimilarity between aboveground vegetation and seed rain increased with species richness in the aboveground vegetation (χ²_Aboveground richness_= 13.6, p < 0.001, Figure 3c). The variation explained by the model including LUI, productivity and species richness in the vegetation as predictors was 38 % (Supplementary material 2).

## Discussion

Our study shows that several key properties of the seed rain of 142 agriculturally managed grasslands were related to land-use intensity, productivity, and aboveground species richness. The total number of seedlings per m² varied between 90 up to ∼ 68,000. This extreme variability is compatible with other studies in grasslands (see e.g., Jakobsson et al., 2006), and can severely complicate deriving patterns from small-scale seed rain studies. Nevertheless, we identified several environmental drivers that directly affect seed rain, indicating that a large sample size and long environmental gradients (such as land-use intensity) is required to explain variation in seed rain.

### Seed rain functional and species composition

We found four times more grass than forb seeds in the seed rain. Their high cover in the aboveground vegetation and their abundance of flowering ramets (Jakobsson et al., 2006; Scotton, 2020) probably causes their dominance in seed rain. The dominance of grasses in seed rain is reinforced by their early phenological development. Martínková et al. (2002) showed that many graminoids start their reproduction cycle earlier than forbs and are usually able to produce ripe seeds before the first cut or the onset grazing at the end of May, whereas many forbs complete their cycle later, during midsummer, after the first management intervention. Similarly, Klinger et al. (2021) found that grasses fruit exactly during the main time of mowing (July) in mountain grasslands. Scotton (2020) concluded that many forb species require higher temperatures for sexual reproduction and as a consequence seeds mature later compared to grass species.

Almost 80 % of the 40 000 seedlings found in our study belonged to only ten species. The most numerous species in the seed rain were also the most abundant species in the aboveground vegetation. Most of these are nutrient demanding, agriculturally important grasses such as *Lolium perenne* and *Alopecurus pratensis* (Mielke & Wohlers, 2019), that are supported in their abundance by fertilization and reseeding, or generally common grassland species such as *Poa pratensis agg.* and *Dactylis glomerata*. According to Grime’s plant strategy framework (Grime, 1974), many grasses are classified as competitors, which are able to maximize growth at high nutrient levels. Despite the relatively small seed production per flowering ramet, they can develop numerous flowering ramets early in the season (Kühn et al. 2004). As tall-growing competitors, they are also favored due to their higher inflorescences, which provide a better autochorous dispersal ability (Soons et al., 2004), as well as increased dispersal potential by mowing and grazing (Klinger et al., 2021). In our study, we found no significant increase in the abundance of competitors with increasing land-use intensity. Nonetheless, the abundance of competitors increased slightly in seed rain and significantly in the aboveground vegetation with increasing productivity (Supplementary material 1, Figure S4). As in our study, biomass measurements were conducted before the first cut, a high biomass may indicate a high proportion of competitors in the aboveground vegetation.

### LUI effects on seed rain

Land-use intensity had a negative effect on total and grass seed densities in seed rain, while increasing aboveground richness led to a slight decrease in total seed density. Total seed densities in seed rain showed unimodal distribution along the land-use intensity gradient and linear negative relation with increasing aboveground richness. The slight increase in total seed densities in seed rain with increasing LUI can be attributed to conditions favoring strategy types with higher seed production such as ruderals, while seed losses due to early or frequent management by cutting and grazing are still low. This relation became negative at LUI levels higher than 1.5, where seed production becomes increasingly impeded by early and more frequent disturbances through mowing and grazing while the ability of clonal spread becomes beneficial (Martínková et al., 2020; Plantureux et al., 2005). This can mainly be attributed to the decrease in grass seed production at LUI levels >1.5, indicating that early and frequent mowing and grazing are particularly harmful to the size of seed rain in graminoids while forbs through their on average later and more flexible seed set (Klinger et al., 2021) were less negatively affected by early management interventions. It can be expected that land-use effects on seed rain are even more pronounced in areas with higher cutting frequencies (up to 6 times per year) and fertilizer input, but even the land-use intensities considered in our study may strongly affect the regeneration of grassland communities.

Beyond a LUI threshold of 1.5, species richness in the seed rain significantly declined, which runs in parallel with a similar decline of species-richness in the aboveground vegetation with increasing LUI (Allan et al., 2014). Our results clearly confirm the assumption that species richness in seed rain is strongly linked to the small-scale local species pool in established vegetation due to the short dispersal distances of many grassland species (Diacon-Bolli et al., 2013; Harper, 1978). This is especially true for low growing species, which are characterized by very short dispersal distances (Soons et al., 2004). Although we did not find a significant effect of aboveground productivity on the species richness in the seed rain, there is a well established negative impact of productivity on species richness in aboveground vegetation (Kleijn et al., 2008; Socher et al., 2012). Thus, we assume an indirect negative effect of productivity on the number of species in seed rain via a reduction of the local aboveground species pool.

We found furthermore a significant increase of the abundance of ruderals in seed rain with increasing LUI. Ruderal plants are adapted to frequent disturbances, reproduce quickly, and rapidly colonize disturbed micro-habitats. Examples of such ruderals in seed rain are the two most abundant forb species, *Cerastium holosteoides* and *Veronica arvensis*. Both species did not have high cover values in aboveground vegetation but occurred at most plots. They start flowering early in March and are characterized by an extended period of flowering (Kühn et al., 2004). Thus, despite their low cover in aboveground vegetation they are able to produce a high number of small seeds and tend to form long-term persistent seed bank (Grime, 1974). Finally, we found decrease in the amount of stress tolerators with increasing land-use intensity. This can be expected, as stress tolerators are characterized by slow growth rate and low investment in reproduction. Further, in our study, the plots with lowest LUI and productivity were mainly sheep-grazed pastures on calcareous limestone soils. These low productive grasslands, where plants must cope with water and severe nutrient limitation, are characterized by the dominance of stress tolerant species (Grime, 1979; Supplementary material 1, Figure S4). However, the low seed production in these nutrient poor grassland may counterbalanced to some degree, as a high proportion of seeds can mature due to late grazing (or mowing) and low livestock densities.

Among the components of land-use intensity (i.e. fertilization, mowing, and grazing), fertilization was the main driver of seed rain density and functional composition. Fertilization increases productivity and causes a shift in plant species composition that favors more competitive species that can transform high nutrient inputs into biomass in the long run (Socher et al., 2012). The altered species composition leads to an increased seed production, mainly by grasses and less of forbs (Zeiter et al. 2013; Scotton & Rossetti 2021). However, higher fertilizer applications are usually accompanied by earlier and more frequent mowing (Blüthgen et al., 2012). While mowing initially had a slight positive effect on species richness and total seed density in the seed rain, higher mowing frequencies (>one cut per year) had negative effects. The positive effect of low cutting frequencies can be attributed to the fact that mowing enhances species-richness in aboveground vegetation by the reduction of litter layers and the creation of temporary gaps that favor the recruitment of low competitive species (Doisy et al., 2014). At higher cutting frequencies, however, seed production is inhibited, as the first cut happens earlier and time between successive cuts decreases. Grazing had a pronounced negative effect on species richness in the seed rain and significantly reduced the seed density of grasses while having no effect on forbs. This may be due to many forbs having evolved physical and chemical defense mechanisms against large ungulate herbivory (Strauss et al., 2002; Veldman et al., 2015). Grasslands with high grazing scores in LUI are usually grazed early in the season before grasses are flowering (Mielke & Wohlers, 2019) and are additionally additionally mown to remove ungrazed plants, which strongly limits seed production (Morris, 2000). Nonetheless, high grazing pressure also increases the availability of open soil gaps in the vegetation that may strongly favor the establishment of forbs following a ruderal life strategy (Bråthen et al., 2021). In fact, ruderal herbs with a high seed production such as *Taraxacum* spec., *Veronica arvensis*, and *Cerastium holosteoides* persisted and even flourished under high grazing pressure in our study, as they are able to produce a high number of seeds early in the season (Grime, 1974; Kühn et al., 2004).

### Land-use effects on dissimilarity between vegetation and seed rain

The Sørensen dissimilarity between vegetation and seed rain was significantly and non-linearly affected by land-use intensity, increasing below and above LUI∼1.5. While at low land-use intensities, dissimilarity was high due to higher aboveground species richness and lower seed production per species, very high land-use intensities led to fewer species producing seeds and higher dispersal distances. Additionally, seed releasing height has been reported to be oppositely affected by different management practices: Intensive grazing promotes the formation of short lawns, where phenotypes of species with low seed releasing height are favored (Plantureux et al., 2005). In contrast, on highly fertilized and frequently cut meadows, the remaining plants may be able to regrow between cuts. Our findings clearly show that dissimilarity between aboveground vegetation and seed rain significantly increased with increasing species-richness in aboveground vegetation. Interestingly, increasing productivity increased the similarity between aboveground vegetation and seed rain only in the Alb region, whereas no relationship was evident in the Schorfheide and Hainich regions. Increasing productivity usually leads to a decrease in species-richness of agricultural grasslands (Klaus et al., 2013; Socher et al., 2012). Additionally, as dominant plants produce more seeds owing to a better nutrient supply (Fujita et al., 2014), the representation of their seeds in our seed rain samples becomes more likely. This is even more true as taller growing vegetation exhibits an elevated seed releasing height that allow seeds to be dispersed over longer distances and more homogeneously (Soons et al., 2004). In conjunction with the spatial uniformity of productive grasslands regarding soil properties (e.g. soil moisture, nutrients) or vegetation structures (e.g. shrubs, fallow areas), this causes a spatial homogenization of plant species assemblages at sites with low aboveground richness (Chisté et al., 2018).

Vice-versa, low productive species-rich sites show in general a much lower seed production per species combined with a higher spatial heterogeneity of microhabitats, which is considerably lowering the probability that seeds of a certain species are represented in our seed traps. This is particularly true for rare species, which may be under-represented in this study, as they mainly shed seeds close to the parent plant (Harper, 1978). This is in line with our finding that dissimilarity between seed rain and aboveground vegetation increases linearly with species-richness.

## Conclusions

Our results demonstrate that the functional and taxonomic diversity of seed rain are negatively affected by land-use intensity. While an increased productivity results in a higher seed density in the seed rain, fewer species are able to contribute to the seed rain at high-productivity sites, as more productive grasslands harbor fewer species in aboveground vegetation. This shows how land-use drives the trajectory of grassland community composition through changing the species contributing to seed rain formation. Based on our findings, it can be concluded that intensively used grasslands are less resilient to environmental changes it terms of seed generation ability. To fully understand the impact of land-use induced alterations in seed rain on grassland stability, resilience and biodiversity, further studies integrating aspects of the formation of the soil seed bank, seedling recruitment and interactions with higher trophical levels are needed.

## Author contributions

NH, TK, VK, and DP conceived of the research idea. SK and MF collected data. YPK and SK analyzed the data, YPK and SK with input from LN and SH wrote the manuscript. All authors commented on the manuscript.

## Data availability statement

This work is based on data elaborated by the projects SADE and Escape II of the Biodiversity Exploratories program (DFG Priority Program 1374). The datasets are publicly available in the Biodiversity Exploratories Information System (http://doi.org/10.17616/R32P9Q). The datasets are listed in the references section.

## Acknowledgements

We thank the managers of the three Exploratories, Dr.Miriam Teuscher, Juliane Vogt, Kirsten Reichel-Jung and Florian Staub, and all former managers for their work in maintaining the plot and project infrastructure; Dr. Victoria Grießmeier for giving support through the central office, Andreas Ostrowski for managing the central data base, and Markus Fischer, Eduard Linsenmair, Dominik Hessenmöller, Ingo Schöning, François Buscot, Ernst-Detlef Schulze, Wolfgang W. Weisser and the late Elisabeth Kalko for their role in setting up the Biodiversity Exploratories project. We thank the administration of the Hainich national park, the UNESCO Biosphere Reserve Swabian Alb and the UNESCO Biosphere Reserve Schorfheide-Chorin as well as all land owners for the excellent collaboration. The work has been partly funded by the DFG Priority Program 1374 “Biodiversity-Exploratories” (Project number 60761519). Field work permits were issued by the responsible state environmental offices of Baden-Württemberg, Thüringen, and Brandenburg.

