## Supplementary material 1 for "Impact of land-use intensity, productivity, and aboveground richness on seed rain in temperate grasslands"

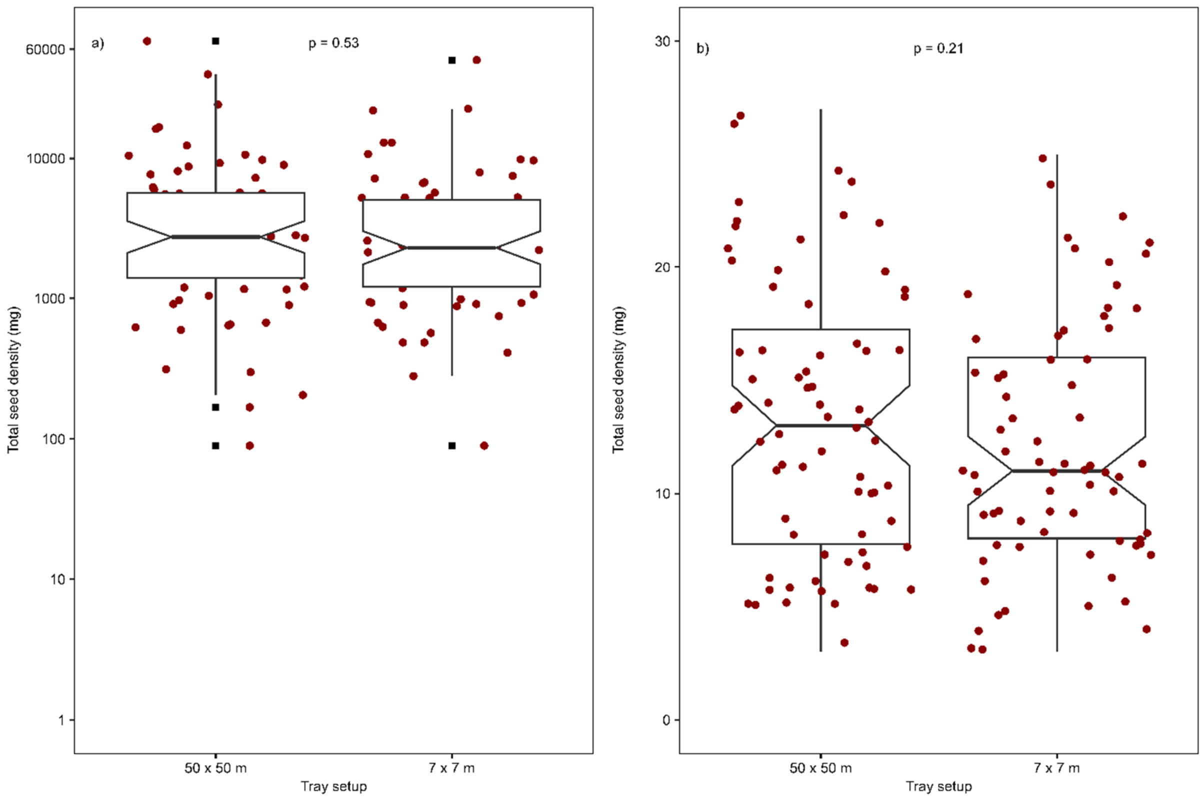


Figure S1: Boxplots comparing both tray setups (50 × 50 m vs. 7 × 7 m) for (a) seed density and (b) aboveground species richness. The upper line of the boxes represents the 75th percentile, the lower line the 25th percentile and the bold middle line the median of the data. The extreme lines represent the highest and lowest value excluding outlier (percentile ± 1.5 × interquartile range). Yellow dots highlight potential outlier in the data. ANOVA was performed to test for significant differences between the tray setups for seed density and aboveground species richness. The tests indicated no significant differences for either seedl density (p value = 0.53) or aboveground species richness (p value = 0.21).

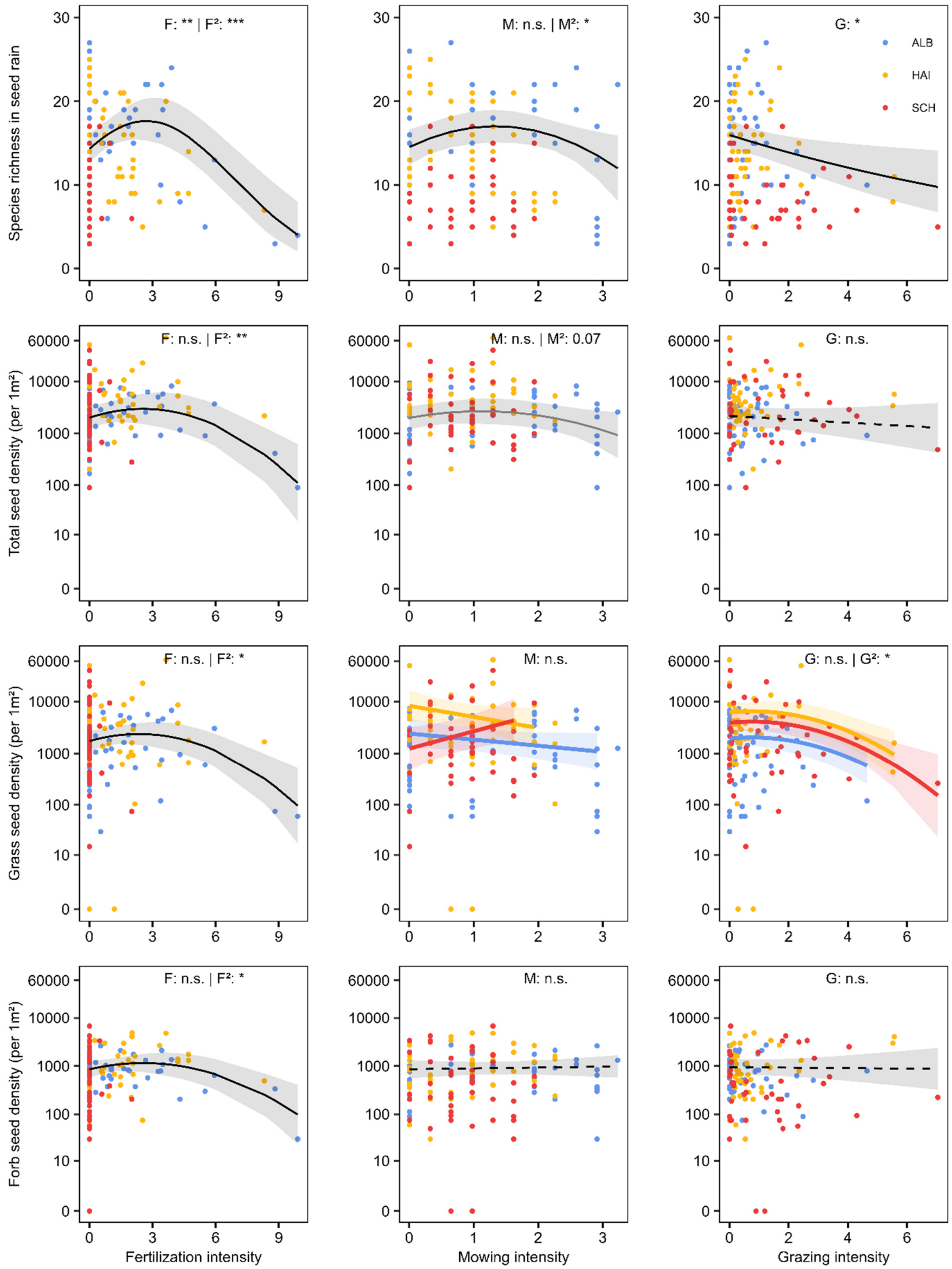


Figure S2: Effects of the single components of LUI (mowing, grazing, fertilizing) and productivity on (a-c) total seed density, the density of the functional groups of (d-f) grasses and (g-i) forbs (legumes included) in seed rain. For (j-l) aboveground species richness, components of LUI and species richness in aboveground vegetation were predictor variables. From (a) to (i) y-axes are log-transformed. Predicted regression lines of the fixed effects based on the mean values of the other predictor variable and region Alb. Grey areas display the confidence interval ($\pm 1.96 \times SE$) of the regression lines. The dots represent the observed data and colours the study regions (Schwäbische Alb (ALB, blue), Hainich-Dün (HAI, orange) and Schorfheide-Chorin (SCH, red). Asterisks indicate significance levels: * $p < 0.05$; ** $p < 0.01$; *** $p < 0.001$ and 'n.s.' = no significance. Marginally non-significant p values are recorded. See Supplementary material 2 for model results.

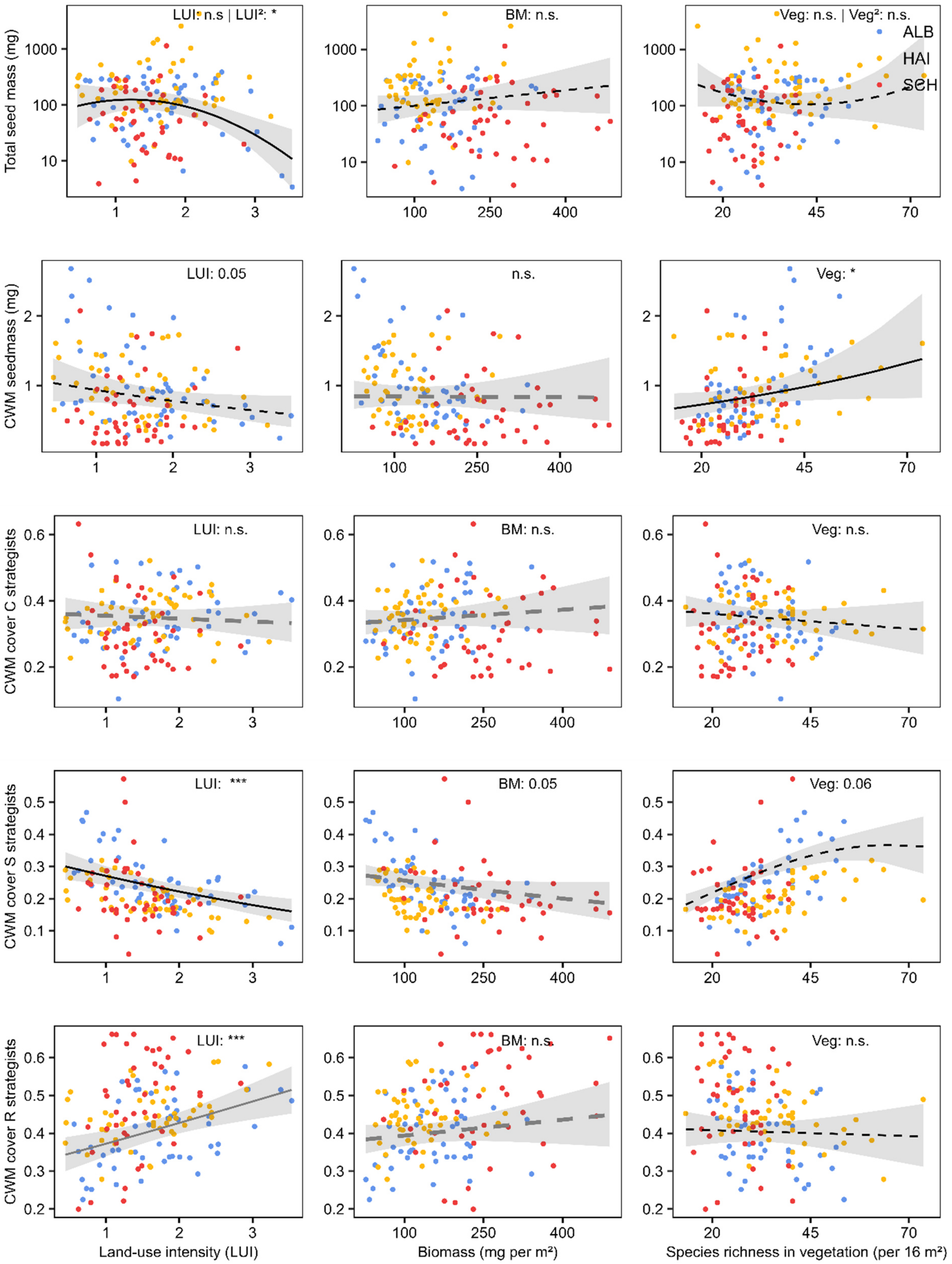


Figure S2: Effects of land-use intensity (LUI), aboveground productivity, and species richness in vegetation on total seed mass in seed rain, community-weighted mean (CWM) seed mass, and cover of C, S, and R strategists in the seed rain. Predicted regression lines of the fixed effects based on the mean values of the other predictor variable and region Alb. Grey areas display the confidence interval (± 1.96 × SE) of the regression lines. The dots represent the calculated values per plot and colors the study regions (Schwäbische Alb (ALB, blue), Hainich-Dün (HAI, orange) and Schorfheide-Chorin (SCH, red). Asterisks indicate significance levels: * p <0.05; ** p <0.01; *** p <0.001 and 'n.s.' = no significance. See Supplementary material 2 for model results. Mind the different scales of y-axes.

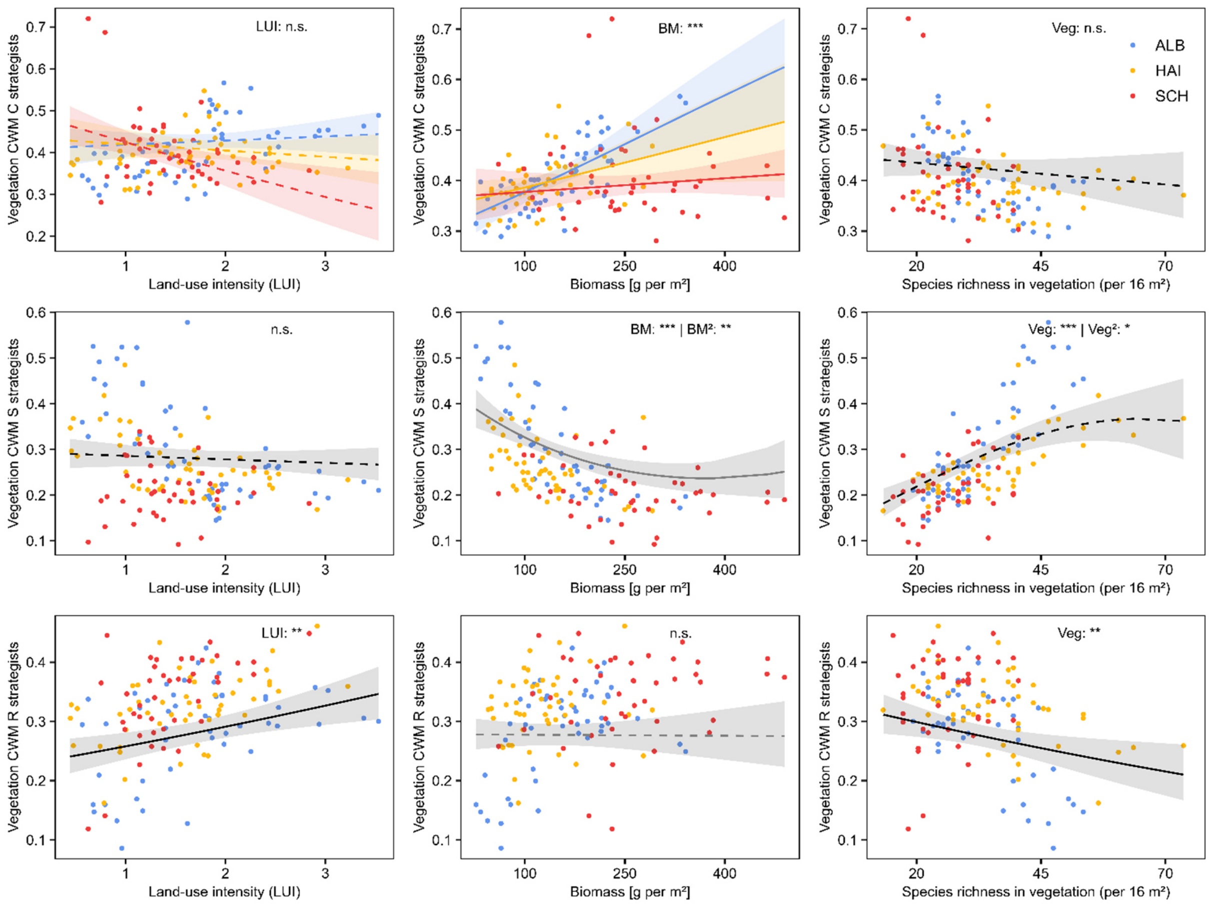


Figure S4: Effects of land-use intensity (LUI), aboveground productivity, and species richness on the cover of C, S, and R strategists in the aboveground vegetation. Predicted regression lines of the fixed effects based on the mean values of the other predictor variable and region Alb. Grey areas display the confidence interval (± 1.96 × SE) of the regression lines. The dots represent the calculated values per plot and colors the study regions (Schwäbische Alb (ALB, blue), Hainich-Dün (HAI, orange) and Schorfheide-Chorin (SCH, red). Asterisks indicate significance levels: * p <0.05; ** p <0.01; *** p <0.001 and 'n.s.' = no significance. See Supplementary material 2 for model results. Mind the different scales of y-axes.
