## Supplementary material 2 for "Impact of land-use intensity, productivity, and aboveground richness on seed rain in temperate grasslands"

**Models containing LUI, Biomass, and aboveground richness**

LUI = Land-use intensity index, calculation according to *Methods* section, consists of Fertilization intensity, Mowing intensity, and Grazing intensity

Biomass= Aboveground productivity averaged between 2016-2018

VEGspecNo= Aboveground species richness averaged between 2016-2018

Explo = Study region

pH_mean = pH measured in $CaCl_2$ solution

twi_full = Topographic wetness index

Organic_C = Content of organic carbon in the topsoil

**Total Seedling density**

Analysis of Deviance Table (Type II tests)

Response: log(SRdensSeedl)

| Variable | LR Chisq | Degrees of Freedom (Df) | P-value (Pr(>Chisq)) | Significance |
|---|---|---|---|---|
| sclLUI1618 | 1.4775 | 1 | 0.224169 | n.s. |
| I(sclLUI1618^2) | 3.616 | 1 | 0.057227 | . |
| sclbiomass1618 | 3.2155 | 1 | 0.072944 | . |
| sclVEGspecNo1618 | 2.9886 | 1 | 0.08385 | . |
| explo | 12.5405 | 2 | 0.001892 | ** |
| pH_mean | 0.0365 | 1 | 0.848449 | n.s. |
| twi_full | 0.2407 | 1 | 0.62373 | n.s |
| Organic_C | 1.8375 | 1 | 0.175246 | n.s |

N= 142, $R^2$ = 0.284421

**Grass Seedling density**

Analysis of Deviance Table (Type II tests)

Response: log10(SRdensGrass + 1)

| Variable | LR Chisq | Degrees of Freedom (Df) | P-value (Pr(>Chisq)) | Significance |
|---|---|---|---|---|
| sclLUI1618 | 0.0793 | 1 | 0.77819 | n.s. |
| I(sclLUI1618^2) | 4.5253 | 1 | 0.0334 | * |
| sclVEGspecNo1618 | 5.8175 | 1 | 0.01587 | * |
| I(sclVEGspecNo1618^2) | 6.1502 | 1 | 0.01314 | * |
| pH_mean | 0.0744 | 1 | 0.78501 | n.s. |
| twi_full | 0.0621 | 1 | 0.80326 | n.s. |
| Organic_C | 3.9208 | 1 | 0.04769 | * |

N= 142, $R^2$ = 0.1145694

**Forb Seedling density**

Analysis of Deviance Table (Type III tests)

Response: ceiling(SRdensDicotyl)

| Variable | LR Chisq | Degrees of Freedom (Df) | P-value (Pr(>Chisq)) | Significance |
|---|---|---|---|---|
| sclLUI1618 | 0.3692 | 1 | 0.54345 | n.s. |
| sclbiomass1618 | 3.8082 | 1 | 0.051002 | . |
| sclVEGspecNo1618 | 3.3147 | 1 | 0.068662 | . |
| I(sclVEGspecNo1618^2) | 8.801 | 1 | 0.003011 | ** |
| explo | 6.557 | 2 | 0.037685 | * |
| pH_mean | 1.0489 | 1 | 0.30577 | n.s. |
| twi_full | 0.297 | 1 | 0.585767 | n.s. |
| Organic_C | 2.1508 | 1 | 0.142497 | n.s. |
| sclLUI1618:explo | 8.6205 | 2 | 0.01343 | * |

N= 142, $R^2$ = 0.2900355

**Species Number in Seed Rain**

Analysis of Deviance Table (Type II tests)

Response: SRspecNo

| Variable | LR Chisq | Degrees of Freedom (Df) | P-value (Pr(>Chisq)) | Significance |
|---|---|---|---|---|
| sclLUI1618 | 0.9352 | 1 | 0.33352 | n.s. |
| I(sclLUI1618^2) | 6.0697 | 1 | 0.01375 | * |
| sclVEGspecNo1618 | 3.9075 | 1 | 0.04807 | * |
| explo | 23.1467 | 2 | 9.41E-06 | *** |
| pH_mean | 2.7407 | 1 | 0.09782 | . |
| twi_full | 0.0266 | 1 | 0.87049 | n.s. |
| Organic_C | 0.0545 | 1 | 0.81549 | n.s. |
| sclbiomass1618 | 0.401 | 1 | 0.52656 | n.s. |

N=142, $R^2$= 0.385

**Total Seedling density and LUI components**

**Fertilization**

Analysis of Deviance Table (Type II tests)

Response: log(SRdensSeedl)

| Variable | LR Chisq | Degrees of Freedom (Df) | P-value (Pr(>Chisq)) | Significance |
|---|---|---|---|---|
| sclF_1618 | 0.6652 | 1 | 0.414737 | |
| I(sclF_1618^2) | 6.8561 | 1 | 0.008834 | ** |
| sclbiomass1618 | 5.3853 | 1 | 0.020308 | * |
| I(sclVEGspecNo1618^2) | 1.4232 | 1 | 0.232885 | |

| | | | | |
|---|---|---|---|---|
| explo | 8.1921 | 2 | 0.016638 | * |
| pH_mean | 0.0255 | 1 | 0.873056 | |
| twi_full | 0.0164 | 1 | 0.897973 | |
| Organic_C | 2.8864 | 1 | 0.089329 | . |
| sclVEGspecNo1618 | 0 | 1 | 0.998772 | |
| sclF_1618:explo | 5.6435 | 2 | 0.059501 | . |
| I(sclF_1618^2):explo | 4.2725 | 2 | 0.118099 | |
| I(sclVEGspecNo1618^2):explo | 4.492 | 2 | 0.105821 | |

N= 142, $R^2$=0.413872

**Mowing**

Analysis of Deviance Table (Type II tests)

Response: log(SRdensSeedl)

| Variable | LR Chisq | Degrees of Freedom (Df) | P-value (Pr(>Chisq)) | Significance |
|---|---|---|---|---|
| sclM_1618 | 0.1161 | 1 | 0.73327 | |
| I(sclM_1618^2) | 3.2572 | 1 | 0.07111 | . |
| sclbiomass1618 | 5.0056 | 1 | 0.02526 | * |
| I(sclVEGspecNo1618^2) | 1.1976 | 1 | 0.2738 | |
| explo | 8.2246 | 2 | 0.01637 | * |
| pH_mean | 0.0533 | 1 | 0.81741 | |
| twi_full | 0.039 | 1 | 0.84339 | |
| Organic_C | 3.3146 | 1 | 0.06867 | . |
| sclVEGspecNo1618 | 0.0037 | 1 | 0.95117 | |
| I(sclVEGspecNo1618^2):explo | 5.5624 | 2 | 0.06196 | . |

N=142, $R^2$=0.2978287

**Fertilization**

Analysis of Deviance Table (Type III tests)

Response: log(SRdensSeedl)

| Variable | LR Chisq | Degrees of Freedom (Df) | P-value (Pr(>Chisq)) | Significance |
|---|---|---|---|---|
| sclG_1618 | 0.8107 | 1 | 0.367926 | |
| sclbiomass1618 | 5.5939 | 1 | 0.018023 | * |
| I(sclVEGspecNo1618^2) | 0.3953 | 1 | 0.529525 | |
| explo | 12.7093 | 2 | 0.001739 | ** |
| pH_mean | 0.0787 | 1 | 0.779001 | |
| twi_full | 0.0251 | 1 | 0.874177 | |
| Organic_C | 2.7699 | 1 | 0.096055 | . |
| sclVEGspecNo1618 | 0.0998 | 1 | 0.752059 | |
| I(sclVEGspecNo1618^2):explo | 5.6875 | 2 | 0.058206 | . |

N=142, $R^2$=0.273086

**Grass Seedling density and LUI components**

**Fertilization**

Analysis of Deviance Table (Type III tests)

Response: SRdensGrass

| Variable | LR Chisq | Degrees of Freedom (Df) | P-value (Pr(>Chisq)) | Significance |
|---|---|---|---|---|
| sclF_1618 | 1.4468 | 1 | 0.2290494 | |
| I(sclF_1618^2) | 4.7595 | 1 | 0.0291366 | * |
| sclbiomass1618 | 2.5004 | 1 | 0.1138206 | |
| sclVEGspecNo1618 | 0.0069 | 1 | 0.9339846 | |
| explo | 15.3983 | 2 | 0.0004532 | *** |
| pH_mean | 0.7456 | 1 | 0.3878674 | |
| twi_full | 0.0006 | 1 | 0.9806473 | |
| Organic_C | 3.493 | 1 | 0.0616275 | . |
| sclVEGspecNo1618:explo | 7.383 | 2 | 0.0249342 | * |

N=142, $R^2$=0.3252702

**Mowing**

Analysis of Deviance Table (Type III tests)

Response: SRdensGrass

| Variable | LR Chisq | Degrees of Freedom (Df) | P-value (Pr(>Chisq)) | Significance |
|---|---|---|---|---|
| sclM_1618 | 1.9176 | 1 | 0.1661257 | |
| sclbiomass1618 | 7.3584 | 1 | 0.0066749 | ** |
| I(sclVEGspecNo1618^2) | 0.2421 | 1 | 0.6226804 | |
| explo | 15.676 | 2 | 0.0003945 | *** |
| pH_mean | 0.0058 | 1 | 0.9390568 | |
| twi_full | 0.0426 | 1 | 0.8365103 | |
| Organic_C | 3.2706 | 1 | 0.0705297 | . |
| sclVEGspecNo1618 | 1.4852 | 1 | 0.2229641 | |
| sclM_1618:explo | 6.0304 | 2 | 0.0490359 | * |
| I(sclVEGspecNo1618^2):explo | 6.0713 | 2 | 0.0480429 | * |

N=142, $R^2$=0.3208993

**Grazing**

Analysis of Deviance Table (Type II tests)

Response: SRdensGrass

| Variable | LR Chisq | Degrees of Freedom (Df) | P-value (Pr(>Chisq)) | Significance |
|---|---|---|---|---|
| sclG_1618 | 0.0438 | 1 | 0.834293 | |
| I(sclG_1618^2) | 4.2194 | 1 | 0.039965 | * |
| sclVEGspecNo1618 | 8.2592 | 1 | 0.004055 | ** |
| explo | 13.5274 | 2 | 0.001155 | ** |
| pH_mean | 0.0398 | 1 | 0.841931 | |
| twi_full | 0.0008 | 1 | 0.97775 | |
| Organic_C | 1.9591 | 1 | 0.161606 | |

N=142, $R^2$=0.2723378

**Forb Seedling density and LUI components**

**Fertilization**

Analysis of Deviance Table (Type III tests)

Response: ceiling(SRdensDicotyl)

| Variable | LR Chisq | Degrees of Freedom (Df) | P-value (Pr(>Chisq)) | Significance |
|---|---|---|---|---|
| sclF_1618 | 1.2549 | 1 | 0.2626171 | |
| I(sclF_1618^2) | 5.9937 | 1 | 0.0143574 | * |
| sclVEGspecNo1618 | 1.7687 | 1 | 0.1835386 | |
| I(sclVEGspecNo1618^2) | 13.0485 | 1 | 0.0003035 | *** |
| explo | 6.6224 | 2 | 0.0364729 | * |
| pH_mean | 3.7799 | 1 | 0.0518713 | . |
| twi_full | 0.4988 | 1 | 0.480042 | |
| Organic_C | 8.0147 | 1 | 0.00464 | ** |
| sclF_1618:explo | 5.2222 | 2 | 0.0734553 | . |
| sclVEGspecNo1618:explo | 6.1558 | 2 | 0.0460551 | * |

N=142, $R^2$=0.3377089

**Mowing**

Analysis of Deviance Table (Type II tests)

Response: ceiling(SRdensDicotyl)

| Variable | LR Chisq | Degrees of Freedom (Df) | P-value (Pr(>Chisq)) | Significance |
|---|---|---|---|---|
| sclM_1618 | 0.1123 | 1 | 0.7375245 | |
| sclVEGspecNo1618 | 0.151 | 1 | 0.6975761 | |
| I(sclVEGspecNo1618^2) | 12.0328 | 1 | 0.0005227 | *** |
| explo | 6.0253 | 2 | 0.0491614 | * |
| pH_mean | 3.0693 | 1 | 0.0797842 | . |

| | | | | |
|---|---|---|---|---|
| twi_full | 1.0259 | 1 | 0.3111195 | |
| Organic_C | 6.9541 | 1 | 0.0083625 | ** |
| sclVEGspecNo1618:explo | 4.5405 | 2 | 0.103286 | |

N=142, $R^2$=0.250329

**Grazing**

Analysis of Deviance Table (Type III tests)

Response: ceiling(SRdensDicotyl)

| Variable | LR Chisq | Degrees of Freedom (Df) | P-value (Pr(>Chisq)) | Significance |
|---|---|---|---|---|
| sclG_1618 | 0.0231 | 1 | 0.879139 | |
| sclbiomass1618 | 2.0791 | 1 | 0.149331 | |
| sclVEGspecNo1618 | 3.3094 | 1 | 0.068887 | . |
| I(sclVEGspecNo1618^2) | 2.5193 | 1 | 0.112458 | |
| explo | 9.6631 | 2 | 0.007974 | ** |
| pH_mean | 2.5455 | 1 | 0.110612 | |
| twi_full | 0.9848 | 1 | 0.321023 | |
| Organic_C | 5.912 | 1 | 0.015038 | * |
| I(sclVEGspecNo1618^2):explo | 6.8713 | 2 | 0.032205 | * |

N=142, $R^2$=0.2738257

**Species Richness in Seed Rain and LUI components**

**Fertilization**

Analysis of Deviance Table (Type III tests)

Response: SRspecNo

| Variable | LR Chisq | Degrees of Freedom (Df) | P-value (Pr(>Chisq)) | Significance |
|---|---|---|---|---|
| sclF_1618 | 7.1526 | 1 | 0.007486 | ** |
| I(sclF_1618^2) | 19.5272 | 1 | 9.92E-06 | *** |
| I(sclbiomass1618^2) | 2.7307 | 1 | 0.098435 | . |
| sclVEGspecNo1618 | 9.4206 | 1 | 0.002146 | ** |
| I(sclVEGspecNo1618^2) | 11.6043 | 1 | 0.000658 | *** |
| explo | 24.8126 | 2 | 4.09E-06 | *** |
| pH_mean | 3.2532 | 1 | 0.071285 | . |
| twi_full | 0.0017 | 1 | 0.967236 | |
| Organic_C | 0.0038 | 1 | 0.950741 | |
| sclVEGspecNo1618:explo | 7.0889 | 2 | 0.028885 | * |

N=142, Nagelkerke's R2=0.720

**Mowing**

Analysis of Deviance Table (Type III tests)

Response: SRspecNo

| Variable | LR Chisq | Degrees of Freedom (Df) | P-value (Pr(>Chisq)) | Significance |
|---|---|---|---|---|
| sclM_1618 | 2.178 | 1 | 0.140039 | |
| I(sclM_1618^2) | 6.261 | 1 | 0.012342 | * |
| sclVEGspecNo1618 | 6.848 | 1 | 0.008872 | ** |
| I(sclVEGspecNo1618^2) | 10.095 | 1 | 0.001486 | ** |
| explo | 39.357 | 2 | 2.84E-09 | *** |
| pH_mean | 3.063 | 1 | 0.080086 | . |
| twi_full | 0.253 | 1 | 0.615254 | |
| Organic_C | 0.514 | 1 | 0.473348 | |
| sclVEGspecNo1618:explo | 9.527 | 2 | 0.008537 | ** |

N=142, $R^2$=0.418

**Grazing**

Analysis of Deviance Table (Type III tests)

Response: SRspecNo

| Variable | LR Chisq | Degrees of Freedom (Df) | P-value (Pr(>Chisq)) | Significance |
|---|---|---|---|---|
| sclG_1618 | 6.3695 | 1 | 0.0116099 | * |
| sclVEGspecNo1618 | 9.027 | 1 | 0.0026602 | ** |
| I(sclVEGspecNo1618^2) | 14.0307 | 1 | 0.0001799 | *** |
| explo | 29.7302 | 2 | 3.50E-07 | *** |
| pH_mean | 3.8669 | 1 | 0.0492485 | * |
| twi_full | 0.0407 | 1 | 0.8400429 | |
| Organic_C | 0.0441 | 1 | 0.8337207 | |
| sclVEGspecNo1618:explo | 5.5977 | 2 | 0.0608796 | . |

Nagelkerke's $R^2$=0.640

**Sorensen dissimilarity between vegetation and seed rain and LUI, biomass and aboveground richness**

Analysis of Deviance Table (Type III tests)

Response: sor

| Variable | LR Chisq | Degrees of Freedom (Df) | P-value (Pr(>Chisq)) | Significance |
|---|---|---|---|---|
| (Intercept) | 0.7834 | 1 | 0.3761042 | |
| sclbiomass1618 | 21.9576 | 1 | 2.79E-06 | *** |
| sclVEGspecNo1618 | 13.5774 | 1 | 0.0002289 | *** |

| | | | | |
|---|---|---|---|---|
| explo | 63.7169 | 2 | 1.46E-14 | *** |
| pH_mean | 0.7879 | 1 | 0.374738 | |
| twi_full | 1.2502 | 1 | 0.2635162 | |
| Organic_C | 0.6656 | 1 | 0.4145809 | |
| sclLUI1618 | 0.046 | 1 | 0.8301296 | |
| I(sclLUI1618^2) | 27.2947 | 1 | 1.75E-07 | *** |

N=142, $R^2$=0.382226

**Functional characteristics of seed rain and LUI, biomass, and aboveground richness**

**Sum seed mass**

Family: gaussian

Link function: identity

Analysis of Deviance Table (Type II tests)

Response: log(`sum seedmass `)

| Variable | LR Chisq | Degrees of Freedom (Df) | P-value (Pr(>Chisq)) | Significance |
|---|---|---|---|---|
| sclLUI1618 | 2.3475 | 1 | 0.12549 | |
| I(sclLUI1618^2) | 5.2472 | 1 | 0.02198 | * |
| sclbiomass1618 | 1.6418 | 1 | 0.20007 | |
| sclVEGspecNo1618 | 1.8669 | 1 | 0.17183 | |
| I(sclVEGspecNo1618^2) | 2.0354 | 1 | 0.15367 | |
| explo | 23.2856 | 2 | 8.78E-06 | *** |
| pH_mean | 1.0173 | 1 | 0.31315 | |
| twi_full | 0.4473 | 1 | 0.5036 | |
| Organic_C | 4.9126 | 1 | 0.02666 | * |

N=142, $R^2$=0.4709777

**CWM Seed mass**

Family: gaussian

Link function: identity

Analysis of Deviance Table (Type II tests)

Response: log(cwm_seeds)

| Variable | LR Chisq | Degrees of Freedom (Df) | P-value (Pr(>Chisq)) | Significance |
|---|---|---|---|---|
| sclLUI1618 | 3.7514 | 1 | 0.052765 | . |
| sclbiomass1618 | 0.0031 | 1 | 0.955289 | |
| sclVEGspecNo1618 | 3.9505 | 1 | 0.046856 | * |
| explo | 9.3071 | 2 | 0.009528 | ** |
| pH_mean | 3.2844 | 1 | 0.06994 | . |
| twi_full | 0.225 | 1 | 0.635221 | |

| | | | | |
|---|---|---|---|---|
| Organic_C | 5.4174 | 1 | 0.019937 | * |

N=142, $R^2$=0.3679657

**Community-weighted means (CWMs) of Grime's strategy types in Seed Rain**

**CWM C strategists**

Family: beta

Link function: logit

Analysis of Deviance Table (Type II Wald chisquare tests)

Response: cwm_c

| Variable | LR Chisq | Degrees of Freedom (Df) | P-value (Pr(>Chisq)) | Significance |
|---|---|---|---|---|
| sclLUI1618 | 0.3149 | 1 | 0.5747 | |
| sclbiomass1618 | 0.7239 | 1 | 0.3949 | |
| sclVEGspecNo1618 | 0.8476 | 1 | 0.3572 | |
| explo | 7.6338 | 2 | 0.022 | * |
| pH_mean | 0.6389 | 1 | 0.4241 | |
| twi_full | 0.5399 | 1 | 0.4625 | |
| Organic_C | 1.7141 | 1 | 0.1905 | |

N=142, $R^2$=0.09218728

**CWM S strategists**

Family: beta

Link function: logit

Analysis of Deviance Table (Type II Wald chisquare tests)

Response: cwm_s

| Variable | LR Chisq | Degrees of Freedom (Df) | P-value (Pr(>Chisq)) | Significance |
|---|---|---|---|---|
| sclLUI1618 | 14.9545 | 1 | 0.0001101 | *** |
| sclbiomass1618 | 3.8206 | 1 | 0.0506258 | . |
| sclVEGspecNo1618 | 3.4503 | 1 | 0.0632402 | . |
| explo | 9.6543 | 2 | 0.0080094 | ** |
| pH_mean | 2.9673 | 1 | 0.0849649 | . |
| twi_full | 3.6454 | 1 | 0.0562249 | . |

| | | | |
|---|---|---|---|
| Organic_C | 1.166 | 1 | 0.2802299 |

N=142, $R^2$=4199539

**CWM R strategists**

Family: beta

Link function: logit

Analysis of Deviance Table (Type II Wald chisquare tests)

Response: cwm_r

| Variable | LR Chisq | Degrees of Freedom (Df) | P-value (Pr(>Chisq)) | Significance |
|---|---|---|---|---|
| sclLUI1618 | 12.5206 | 1 | 0.0004025 | *** |
| sclbiomass1618 | 1.3546 | 1 | 0.2444813 | |
| sclVEGspecNo1618 | 0.0974 | 1 | 0.7549894 | |
| explo | 9.7138 | 2 | 0.0077746 | ** |
| pH_mean | 4.5547 | 1 | 0.0328279 | * |
| twi_full | 0.275 | 1 | 0.6000117 | |
| Organic_C | 0.7886 | 1 | 0.3745177 | |

N=142, $R^2$=0.2668757

**Community-weighted means (CWMs) of Grime's strategy types in aboveground vegetation**

**Aboveground CWM C strategists**

Family: beta

Link function: logit

Analysis of Deviance Table (Type III Wald chisquare tests)

Response: veg_cwm_c

| Variable | LR Chisq | Degrees of Freedom (Df) | P-value (Pr(>Chisq)) | Significance |
|---|---|---|---|---|
| (Intercept) | 55.0329 | 1 | 1.19E-13 | *** |
| sclLUI1618 | 0.4566 | 1 | 0.4992 | |
| sclbiomass1618 | 16.065 | 1 | 6.12E-05 | *** |
| sclVEGspecNo1618 | 1.2425 | 1 | 0.26499 | |
| explo | 6.6335 | 2 | 0.03627 | * |

| | | | | |
|---|---|---|---|---|
| pH_mean | 1.8508 | 1 | 0.17369 | |
| twi_full | 1.636 | 1 | 0.20087 | |
| Organic_C | 0.1627 | 1 | 0.6867 | |
| sclLUI1618:explo | 8.5244 | 2 | 0.01409 | * |
| sclbiomass1618:explo | 8.3825 | 2 | 0.01513 | * |

N=142, $R^2$=0.3643445

**Aboveground CWM S strategists**

Family: beta

Link function: logit

Analysis of Deviance Table (Type II Wald chisquare tests)

Response: veg_cwm_s

| Variable | LR Chisq | Degrees of Freedom (Df) | P-value (Pr(>Chisq)) | Significance |
|---|---|---|---|---|
| sclLUI1618 | 0.6398 | 1 | 0.42377 | |
| sclbiomass1618 | 25.2038 | 1 | 5.16E-07 | *** |
| I(sclbiomass1618^2) | 7.5257 | 1 | 0.006083 | ** |
| sclVEGspecNo1618 | 36.9898 | 1 | 1.19E-09 | *** |
| I(sclVEGspecNo1618^2) | 6.1843 | 1 | 0.012889 | * |
| explo | 21.7572 | 2 | 1.89E-05 | *** |
| pH_mean | 0.0952 | 1 | 0.75765 | |
| twi_full | 4.9718 | 1 | 0.025764 | * |
| Organic_C | 0.0232 | 1 | 0.87892 | |

N=142, $R^2$=0.7079767

**Aboveground CWM R strategists**

Family: beta

Link function: logit

Analysis of Deviance Table (Type II Wald chisquare tests)

Response: veg_cwm_r

| Variable | LR Chisq | Degrees of Freedom (Df) | P-value (Pr(>Chisq)) | Significance |
|---|---|---|---|---|
| sclLUI1618 | 10.652 | 1 | 0.0011 | ** |

| | | | | |
|---|---|---|---|---|
| sclbiomass1618 | 0.0059 | 1 | 0.938907 | |
| sclVEGspecNo1618 | 6.9812 | 1 | 0.008237 | ** |
| explo | 20.3933 | 2 | 3.73E-05 | *** |
| pH_mean | 4.9979 | 1 | 0.025378 | * |
| twi_full | 0.9167 | 1 | 0.338349 | |
| Organic_C | 1.8728 | 1 | 0.171151 | |

N=142, $R^2$=0.341358
